# APOLIPOPROTEIN-E (APOE) ε4 AND COGNITIVE DECLINE OVER THE ADULT LIFE COURSE

**DOI:** 10.1101/207233

**Authors:** Mark James Rawle, Daniel Davis, Rebecca Bendayan, Andrew Wong, Diana Kuh, Marcus Richards

## Abstract

We tested the association between APOE-ε4 and processing speed and memory between ages 43 and 69 in a population-based birth cohort. Analyses of processing speed (using a timed letter search task) and episodic memory (a 15-item word learning test) were conducted at ages 43, 53, 60-64 and 69 years using linear and multivariable regression, adjusting for gender and childhood cognition. Linear mixed models, with random intercepts and slopes, were conducted to test the association between APOE and the rate of decline in these cognitive scores from age 43 to 69. Model fit was assessed with the Bayesian Information Criterion. A cross-sectional association between APOE-ε4 and memory scores was detected at age 69 for both heterozygotes and homozygotes (β=−0.68 & β=−1.38 respectively, p=.03) with stronger associations in homozygotes; no associations were observed before this age. Homozygous carriers of APOE-ε4 had a faster rate of decline in memory between ages 43 and 69, when compared to noncarriers, after adjusting for gender and childhood cognition (β=−0.05, p=.04). There were no cross-sectional or longitudinal associations between APOE-ε4 and processing speed. We conclude that APOE-ε4 is associated with a subtly faster rate of memory decline from midlife to early old age; this may be due to effects of APOE-ε4 becoming manifest around the latter stage of life. Continuing follow-up will determine what proportion of this increase will become clinically significant.

## INTRODUCTION

Apolipoprotein E (APOE) is involved in the transport of cholesterol and other lipids between cellular structures.^1^ It is genetically associated with two single nucleotide polymorphisms (SNPs) which mark three alleles, ε2, ε3 and ε4. The ε4 in particular has a higher rate of lipoprotein clearance thus altering both plasma cholesterol level and disrupting brain re-innervation processes that rely on lipids.^2^ APOE is also involved in clearing beta amyloid from the brain, and the ε4 allele may be less efficient at this.^3^ The combined effect of changes in these mechanisms means individuals heterozygous for the ε4 genotype of APOE have three times the risk of developing sporadic Alzheimer’s Disease (AD), increasing to fourteen times the risk for homozygotes.^2,4^

Associations between APOE-ε4 and incident AD have been demonstrated in numerous epidemiological studies,^2,4–6^ yet the effects of APOE-ε4 on decline across the full spectrum of cognitive capability in the general population are less clear. A multi-cohort consortium that included the Medical Research Council National Survey for Health and Development (NSHD), found that APOE genotype had no significant effects on multiple cognitive and functional domains across midlife and old age,^7^ consistent with other studies examining cognitive trajectories in younger individuals.^8–10^ However a comprehensive meta-analysis of the effects of APOE on cognition across a wider range of age groups in population-based cohorts suggested that the ε4 allele has a small but significant negative effect on cognition, more pronounced in older age groups.^11^ While APOE associated cognitive has been noted within birth cohort studies in the ninth decade,^12^ a longitudinal analysis conducted on a mixed age group sample of known APOE status additionally noted an increased rate of decline in memory in homozygous ε4 carriers, manifesting prior to age 60.^13^ Yet with a follow up of five years, and age heterogeneity among the sample population, further work to support or refute the findings of this study would be valuable.

Since its inclusion in the above meta-analysis,^7^ the NSHD has undergone two further waves of assessment, providing a total of four repeated measures of verbal episodic memory and processing speed spanning ages 43 and 69 years.^14^ This birth cohort offers two strong advantages for investigating the associations between APOE genotype and later-life cognition: age homogeneity and individual cognitive trajectories available from midlife (when age-associated cognitive impairment is minimal) to early old age (when this is more pronounced but frank dementia is still rare). This paper aims to determine the association between APOE-ε4 and processing speed and episodic memory scores from age 43 to 69 in this population-based birth cohort, accounting for childhood cognitive ability.

## METHODS

The MRC National Survey of Health and Development (NSHD) is a birth cohort stratified by paternal social class, focusing on multiple socio-demographic and health related processes across the life course. The initial study consisted of 5362 individuals born in England, Scotland and Wales in a single week of March 1946.^15,16^ Response rates have remained consistently high during the study’s duration, and the sample has remained representative of this generation.^17^

The sample for the current analysis is drawn from age 43, when 3749 participants remained in NSHD. From this group, 3456 (92%) had at least one assessment of cognition at either 43, 53, 60-64 or 69. APOE genotype was available for 2637 of the original study members. A further 67 with the ε2/ε4 genotype were excluded, as associations are difficult to interpret due to opposing effects of these alleles.^18^ Of the individuals with valid APOE data, 2557 (97%) had at least one valid cognitive measure between ages 43 and 69, and 2365 (90%) had data for all co-variables. None of these individuals had a confirmed diagnosis of dementia.

With regard to longitudinal modelling, at least one measure of episodic memory was available for 2352 individuals, with 100 having data from only one time point, 441 from two time points, 498 from three time points, and 1313 from all four time points. For the 2363 individuals with a measure of processing speed 72 had data from a single time point, 437 from two, 495 from three, and 1359 from all four time points.

### Outcome: Adult episodic memory and processing/search speed

Cognitive function was tested at ages 43, 53, 60-64 and 69. Memory was assessed by a 3-trial 15-item word learning test (WLT, max. 45), with word lists alternated over waves to minimise practice effects. Processing speed was assessed by a timed letter search task (TLST), in which study members were required to cross out randomly distributed letters ‘P’ and ‘W’ in a grid of other letters as quickly and accurately as possible in one minute, with a maximum score of 600.

### APOE Genotype

Blood samples were collected at age 53 by a trained research nurse, and DNA was extracted.^19^ Genotyping of the two SNPs, rs439358 and rs7412, used to determine APOE genotype was conducted at LGC, Huddleston UK. For analysis, APOE genotype was recoded for the homozygous or heterozygous presence of ε4 alleles; ε4 carrier was defined as ε3/ε4 or ε4/ε4, with ε2/ε4 excluded. Carriers of ε2 were grouped alongside ‘non APOE-ε4 carriers’, as independent analysis of these ε2 carriers yielded no significant results (likely due to the relative low number of ε2 homozygotes within the sample). Thus APOE was categorised as ‘No APOE-ε4’, ‘Homozygous APOE-ε4’ and ‘Heterozygous APOE-ε4’.

### Covariables

Covariables were factors known to be associated with cognitive performance in later life: gender, childhood cognitive ability, formal educational qualifications^20^ and the presence of vascular risk factors that might potentially have a role in the association between APOE and cognition.^21^

Childhood cognitive ability at age 8 was represented as the standardised sum of four tests of verbal and non-verbal ability devised by the National Foundation for Educational Research that included reading comprehension, pronunciation, vocabulary and nonverbal reasoning.^22^ Data were used from age 8 unless missing, where it was substituted for z-scores from the assessments at age 11 (n=128) Data still missing were substituted for that collected at age 15 (n=71). Test-retest correlation coefficients indicated high reliability for each of these standardised scores.^22,23^

Highest educational qualifications by age 26 were classified using the Burnham Scale^24^ and were grouped into three levels for analysis: ‘None’, ‘Vocational or O-Level/GCSE or equivalents’, ‘A-Level or Higher Education or equivalents’. Social class was recorded at age 53, or earlier if this was missing, based on the Registrar General’s classification of own occupation. Vascular risk factors were represented by clinical disorders at age 60 to 64, summarised categorically as the presence of moderate to severe diabetes, hypertension, obesity, hypercholesterolaemia, or cerebrovascular and cardiovascular disease.^25^ Measures of total cholesterol were available at ages 53, 60-64 and 69 recorded in millimoles per litre to the nearest decimal place.

#### Ethics

For the most recent data collection, ethical approval was obtained from the NRES Queen Square REC (14/LO/1073) and Scotland A REC (14/SS/1009). At each stage of data collection, written informed consent was provided by all participating study members. Ethical approval of NSHD data collection up to 2010 was obtained from the Scotland A Research Ethics Committee, the Multicentre Research Ethics Committee, and consent for the use of genetic data in studies of health was obtained from the Central Manchester Research Ethics Committee.

#### Statistical Methods

Differences in social class, educational qualifications and childhood cognitive ability by APOE genotype were assessed using chi-square and t-tests. Associations between APOE and each cognitive test score were analysed using linear and multivariate regression models separately for each of the four time points. Adjustments were initially made sequentially for gender, childhood cognition, education and vascular risk factors.

Linear mixed models were used to investigate the trajectories of verbal memory and search speed scores and their association with APOE. Intercepts and slopes were random to account for inter-individual variability at baseline and the rate of change. Linear and quadratic models were examined and an unstructured covariance structure was assumed. Associations with intercept and slope were sequentially estimated for gender and childhood cognitive ability, education and vascular risk factors. Model fit was compared by using the Bayesian Information Criterion (BIC). First, unconditional linear and quadratic models were compared and then conditional models were estimated. These were as follows; *Model 1*: APOE. *Model 2*: adjusted for gender. *Model 3*: additionally adjusted for childhood cognitive ability. *Model 4*: additionally adjusted for education. *Model 5*: additionally adjusted for vascular risk factors. Analysis was initially conducted on cases with data available for all covariates. Where the inclusion of covariates were found to have no effect on associations between APOE and cognition, the covariate was removed from the final model and those persons omitted only due to missing data from this now removed co-variate were reincorporated. Statistical software package Stata version 14.1 was used for all analyses.

## RESULTS

The APOE genotype frequencies within NSHD were as follows, ε2/ε2 n=20 (0.76%), ε2/ε3 n=307 (11.64%), ε3/ε3 n=1520 (57.64%), ε2/ε4 n=67 (2.54%), ε3/ε4 n=639 (24.23%), ε4/ε4 n=84 (3.19%). Sample characteristics for all individuals with recorded APOE status are outlined in Table 1.

**TABLE 1:** SAMPLE CHARACTERISTICS (BY APOE-ε4 STATUS)

|  | No APOE-ε4 | Heterozygous APOE-ε4 | Homozygous APOE-ε4 | Missing |
| --- | --- | --- | --- | --- |
| **Sample Size** | 1699 (71.8%) | 590 (25.0%) | 76 (3.2%) | 2997 (55.9%) |
| **Female** | 953 (51.6%) | 293 (45.9%) | 45 (53.6%) | 522 (47.9%) |
|  |  |  | *p*= .04 | *p*< .01 |
| **Childhood Cognition\* μ (s.d.)** | 0 (1.0) | 0 (1.0) | 0.1 (1.0) | 0 (1.0) |
| **Education Status age 26:** |  |  |  |  |
| None | 636 (36.5%) | 211 (34.8%) | 31 (37.8%) | 932 (43.7%) |
| Vocational / O-Level | 491 (28.2%) | 170 (28.0%) | 23 (28.1%) | 567 (26.6%) |
| A-Level / Higher | 617 (35.4%) | 226 (37.2%) | 28 (34.2%) | 635 (29.8%) |
|  |  |  | *p*= .92 | *p*< .01 |
| **Social Class age 53:** |  |  |  |  |
| Unskilled | 82 (4.5%) | 24 (3.8%) | 1 (1.2%) | 76 (4.1%) |
| Partly Skilled | 209 (11.4%) | 80 (12.5%) | 11 (13.1%) | 260 (14.1%) |
| Skilled Manual | 321 (17.6%) | 110 (17.2%) | 15 (17.9%) | 401 (21.8%) |
| Skilled Non-Manual | 424 (23.2%) | 141 (22.1%) | 16 (19.1%) | 445 (24.1%) |
| Intermediate | 674 (36.9%) | 226 (35.4%) | 35 (41.7%) | 542 (29.4%) |
| Professional | 118 (6.5%) | 57 (8.9%) | 6 (7.1%) | 120 (6.5%) |
|  |  |  | *p*= .55 | *p*< .01 |
| **Clinical Disorders 60-64:** |  |  |  |  |
| Hypercholesterolemia | 230 (23.1%) | 111 (31.3%) | 12 (27.3%) | 77 (23.7%) |
|  |  |  | *p*= .01 | *p*= .48 |
| Hypertension | 701 (52.9%) | 238 (52.1%) | 28 (47.5%) | 260 (52.7%) |
|  |  |  | *p*= .70 | *p*= .94 |
| Cardio/Cerebrovascular | 108 (8.6%) | 42 (12.2%) | 7 (13.5%) | 45 (8.4%) |
|  |  |  | *p*= .06 | *p*= .30 |
| Diabetes | 254 (23.9%) | 95 (25.2%) | 10 (21.3%) | 87 (24.4%) |
|  |  |  | *p*= .79 | *p*= .97 |
| Obesity | 396 (29.7%) | 128 (28.0%) | 17 (28.3%) | 141 (28.5%) |
|  |  |  | *p*= .76 | *p*= .64 |

\*Derived from Childhood Cognitive Testing Battery, Standardised Value using mean=1, Standard Deviation =0.

Childhood cognitive ability, despite having no association with APOE genotype, was still included in all models, since this allows each individual to begin from an equal cognitive level in the early life course, thus allowing more precise assessment of the longitudinal effects of APOE genotype on cognition. Of those with known APOE status, 2365 (92%) had a valid measure of childhood cognitive ability. Education bore no association with APOE genotype and had no impact on associations between APOE genotype and cognition in either cross sectional or longitudinal models, and was consequently dropped from final models. Sensitivity analyses were conducted on measured vascular risk factors, and the inclusion of all vascular risk factors did not alter any associations seen between APOE and cognition. Only the presence of hypercholesterolaemia displayed variation between APOE genotypes. Therefore an additional sensitivity analysis of cholesterol levels alone, available for ages 53, 60-64 and 69, was conducted on cross sectional models at these ages, but these too had no impact upon the association of APOE genotype with cognition. Additionally, the absence of a cholesterol measurement at age 43 prevented its inclusion in the longitudinal model of cognition, and thus it was omitted from the final models. As such, the final sample consisted of individuals with data on APOE genotype and childhood and adult cognitive scores. Sensitivity analyses showed no differences in the results when missing childhood cognitive scores at age 8 were substituted for cognitive scores at age 11 and 15.

### APOE and episodic memory and processing/search speed at age 43, 53, 60-64 and 69

APOE-ε4 was inversely associated with WLT at age 69 only (Table 2). This persisted after adjusting for gender and childhood cognitive ability in both heterozygotes and homozygotes (β=−0.68 and β=−1.38 respectively, p=.03), with a stronger association seen in homozygotes. At ages 43, 53 and 60-64 there were no associations between APOE-ε4 and the memory scores, and no associations between APOE and search speed scores at any age (supplemental table S1).

**TABLE 2:** CROSS-SECTIONAL RESULTS OF APOE STATUS ON TOTAL WORD LEARNING TEST SCORE

|  | Model One (APOE only) |  |  |  | Model Two (APOE + Gender) |  |  |  | Model Three (APOE + Gender + Childhood Cognition) |  |  |  |
| --- | --- | --- | --- | --- | --- | --- | --- | --- | --- | --- | --- | --- |
| | $\beta$ | LCI | UCI | p | $\beta$ | LCI | UCI | p | $\beta$ | LCI | UCI | p |
| **AGE 43 (n=2123)** |  |  |  |  |  |  |  |  |  |  |  |  |
| *No APOE-ε4* | Reference |  |  | .61 | Reference |  |  | .53 | Reference |  |  | .98 |
| *Heterozygous APOE-ε4* | 0.07 | -0.56 | 0.69 |  | 0.15 | -0.47 | 0.77 |  | 0.00 | -0.55 | 0.55 |  |
| *Homozygous APOE-ε4* | 0.77 | -0.76 | 2.30 |  | 0.82 | -0.70 | 2.34 |  | 0.14 | -1.22 | 1.49 |  |
| *BIC* | 13860.78 |  |  |  | 13833.91 |  |  |  | 13351.80 |  |  |  |
| **AGE 53 (n=2313)** |  |  |  |  |  |  |  |  |  |  |  |  |
| *No APOE-ε4* | Reference |  |  | .99 | Reference |  |  | .89 | Reference |  |  | .84 |
| *Heterozygous APOE-ε4* | 0.05 | -0.55 | 0.64 |  | 0.15 | -0.44 | 0.74 |  | 0.01 | -0.50 | 0.53 |  |
| *Homozygous APOE-ε4* | 0.05 | -1.40 | 1.51 |  | 0.08 | -1.36 | 1.51 |  | -0.38 | -1.64 | 0.88 |  |
| *BIC* | 15079.15 |  |  |  | 15039.00 |  |  |  | 14443.10 |  |  |  |
| **AGE 60-64 (n=1672)** |  |  |  |  |  |  |  |  |  |  |  |  |
| *No APOE-ε4* | Reference |  |  | .74 | Reference |  |  | .45 | Reference |  |  | .47 |
| *Heterozygous APOE-ε4* | 0.27 | -0.41 | 0.95 |  | 0.42 | -0.24 | 1.10 |  | 0.33 | -0.27 | 0.92 |  |
| *Homozygous APOE-ε4* | 0.16 | -1.52 | 1.84 |  | 0.21 | -1.45 | 1.87 |  | -0.37 | -1.85 | 1.11 |  |
| *BIC* | 10785.73 |  |  |  | 10746.18 |  |  |  | 10367.02 |  |  |  |
| **AGE 69 (n=1616)** |  |  |  |  |  |  |  |  |  |  |  |  |
| *No APOE-ε4* | Reference |  |  | .14 | Reference |  |  | .21 | Reference |  |  | .03 |
| *Heterozygous APOE-ε4* | -0.64 | -1.32 | 0.05 |  | -0.56 | -1.24 | 0.13 |  | -0.68 | -1.29 | 0.03 |  |
| *Homozygous APOE-ε4* | -0.84 | -2.53 | 0.86 |  | -0.79 | -2.47 | 0.89 |  | -1.38 | -2.89 | 0.07 |  |
| *BIC* | 10443.28 |  |  |  | 10415.69 |  |  |  | 10077.33 |  |  |  |

*All figures rounded to 2 decimal places*

### APOE and Cognitive Decline

BIC statistics showed that a quadratic model better described the trajectories of WLT and TLST from age 43 to 69 than a linear model. For TLST, APOE was not associated with intercept or slope (supplemental table S2).

For WLT, APOE was not significantly associated with intercept, but those homozygous for APOE4 showed a faster linear rate of decline in WLT (β=−0.05, p=.04) when compared with individuals with no APOE. Although a similar trend was found for heterozygotes, the rate of change in this group was not significantly faster when compared to non-carriers (β=−0.01, p=.22). Although gender and childhood cognitive ability were positively associated with WLT at intercept level, no associations were found between these and slope.

Evidence of longitudinal WLT trajectories with regards APOE genotype is provided in Table 3. Figure 1 depicts the cognitive trajectories of WLT mean scores by APOE genotype, with the intercept adjusted for gender and childhood cognitive ability. This indicates a faster rate of cognitive decline for those individuals homozygous for APOE-ε4.

**FIGURE 1:**
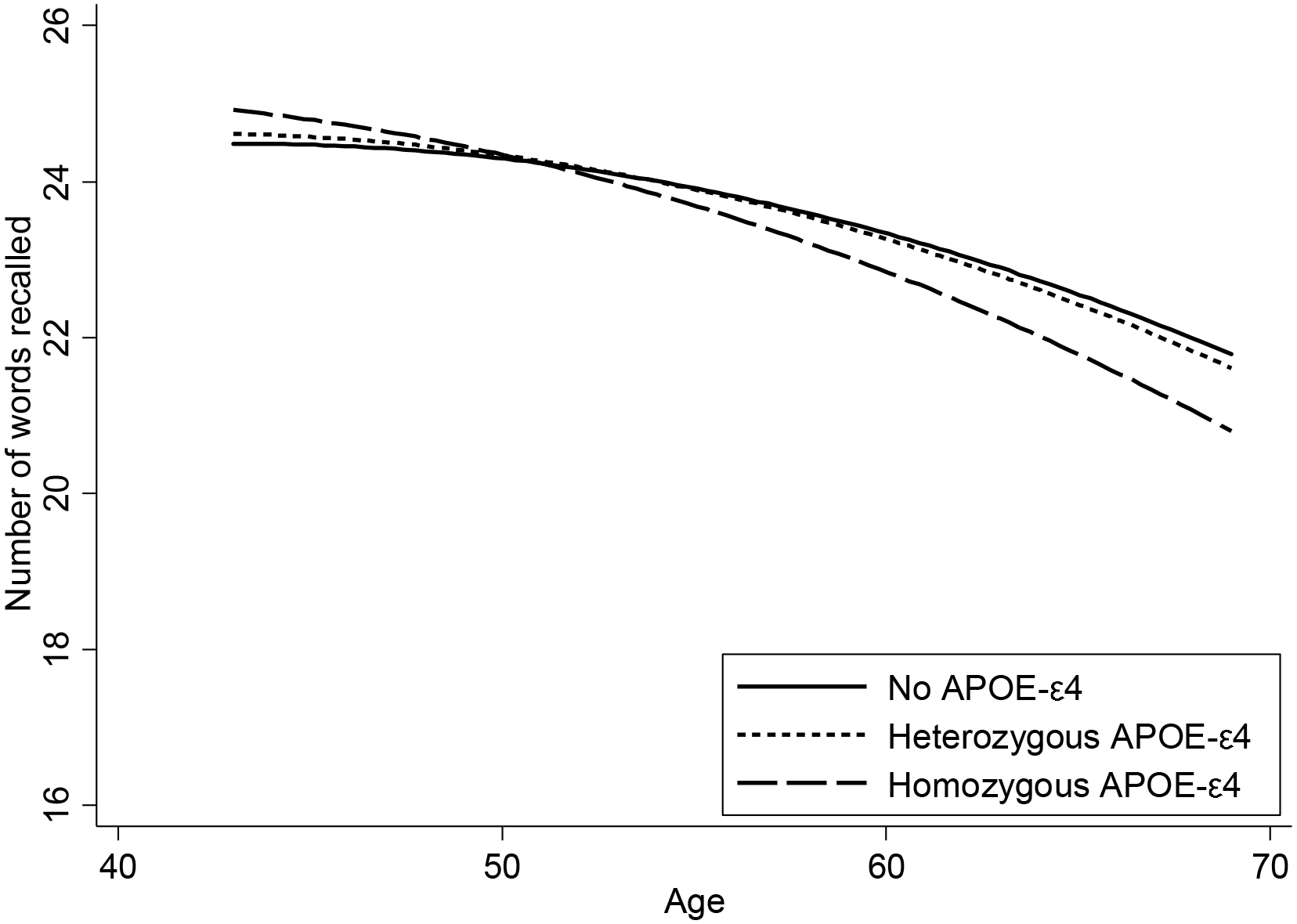
COGNITIVE TRAJECTORY OF TOTAL WORD LEARNING TEST SCORE BETWEEN AGES 43 AND 69

**TABLE 3:** LONGITUDINAL RESULTS OF APOE STATUS ON TOTAL WORD LEARNING TEST SCORE BETWEEN AGES 43 AND 69

|  | Model One (APOE only) |  |  |  | Model Two (APOE + Gender) |  |  |  | Model Three (APOE + Gender + Childhood Cognition) |  |  |  |
| --- | --- | --- | --- | --- | --- | --- | --- | --- | --- | --- | --- | --- |
| *INTERCEPT TERMS* | $\beta$ | LCI | UCI | p | $\beta$ | LCI | UCI | p | $\beta$ | LCI | UCI | p |
| APOE-ε4 Status: | | | | | | | | | | | | |
| No APOE-ε4 | Reference | | | | Reference | | | | Reference | | | |
| Heterozygous APOE-ε4 | 0.74 | -0.54 | 2.02 | .26 | 0.84 | -0.44 | 2.11 | .20 | 0.69 | -0.56 | 1.93 | .28 |
| Homozygous APOE-ε4 | 2.72 | -0.37 | 5.81 | .08 | 2.76 | -0.32 | 5.85 | .08 | 2.34 | -0.66 | 5.33 | .13 |
| Gender (Female) |  |  |  |  | 1.82 | 1.37 | 2.27 | <.01 | 1.76 | 1.38 | 2.14 | <.01 |
| Childhood Cognition (8) |  |  |  |  |  |  |  |  | 3.51 | 3.28 | 3.74 | <.01 |
| *SLOPE TERMS* |  |  |  |  |  |  |  |  |  |  |  |  |
| Decline per Year (Linear) | 0.34 | 0.25 | 0.47 | <.01 | 0.34 | 0.21 | 0.47 | <.01 | 0.35 | 0.22 | 0.47 | <.01 |
| Decline per Year (Quadratic) | 0.00 | 0.00 | 0.00 | <.01 | 0.00 | 0.00 | -0.00 | <.01 | 0.00 | 0.00 | 0.00 | <.01 |
| APOE-ε4 Status: | | | | | | | | | | | | |
| No APOE-ε4 Slope | Reference | | | | Reference | | | | Reference | | | |
| Heterozygous APOE-ε4 Slope | -0.01 | -0.03 | 0.01 | .23 | -0.01 | -0.03 | 0.01 | .23 | -0.01 | -0.03 | 0.01 | .22 |
| Homozygous APOE-ε4 Slope | -0.05 | -0.11 | 0.00 | .04 | -0.05 | -0.11 | 0.00 | .04 | -0.05 | -0.11 | 0.00 | .04 |
| BIC | 45268.92 |  |  |  | 45214.72 |  |  |  | 44467.03 |  |  |  |

*All figures rounded to 2 decimal places*

## DISCUSSION

We found that APOE-ε4 had a negative association with verbal episodic memory, as measured by a word-learning test, at age 69. No cross-sectional association was found at any age prior to this, and no association was noted for processing speed as measured by a timed letter search task. Longitudinally, there was evidence of faster decline in episodic memory between ages 43 and 69 in individuals homozygous for APOE-ε4. Taken together, our findings suggest that APOE-ε4 has a dose dependent cumulatively detrimental effect on episodic memory, which becomes evident as a cross-sectional association by age 69.

The major strength of NSHD arises through it being a population-based birth cohort with prospective measures of cognition across life. To our knowledge, this is the first study to investigate the association of APOE and cognitive trajectories over a period of nearly thirty years, also accounting for childhood cognitive ability. Limitations include sample attrition due to participant loss to follow up, inherent to all studies of aging populations.^26^ However, while not random with respect to cognitive function, this is unlikely to have altered the pattern of associations observed. Previous papers have suggested modest associations between vascular risk factors, APOE and cognition;^27,28^ yet within this sample only cholesterol showed associations with APOE genotype. As cholesterol was not measured in NSHD until age 53, it could not be included in the longitudinal model from age 43 onward. However, given the relatively low levels of hypercholesterolemia that might be expected at age 43, it is unlikely to have influenced associations between cognition and APOE-ε4.

The lack of association between APOE genotype and childhood cognition is supportive of prior findings by Deary *et al.*^29^ Consistent with prior work on the NSHD at age 53,^7^ no evidence of APOE associated decline was found in either processing speed or memory at ages 43, 53 and 60-64. The emergence of significantly lower memory between the ages of 60-69 years in APOE-ε4 carriers is consistent with prior cross-sectional and change-based studies,^30–32^ yet with the availability of multiple previous cognitive measures, NSHD uniquely pinpointed this age range as sensitive in this respect. This suggests the possibility of a time window in which individuals at a higher risk of cognitive decline can potentially be identified for future clinical and public health interventions.

Key to this study, the trajectory of episodic memory over the near three decades of testing was also found to show faster decline associated with APOE-ε4 homozygotes. These findings support those of Caselli *et al.*,^13^ who noted an increased rate of memory decline prior to age 60 in homozygotic carriers of the ε4 allele, albeit over a shorter time period in a smaller, age heterogeneous sample. The absence of any association with letter search is similarly consistent with Caselli *et al.* in parallel work on the same cohort.^33,34^

The presence of detectable deficits in episodic memory at age 69 only for APOE-ε4 carriers, coupled with a faster rate of decline leading up to this stage highlights associations that had not been noticeable at earlier data collections of NSHD.^7^ At the cellular level this may be due to disrupted lipid metabolism resulting in reduced ability to perform neural re-innervation,^2^ and a reduced ability to clear neurotoxic beta amyloid.^3^ Recent advances in tau-imaging also suggest the possibility that APOE-ε4 is associated with enhanced tau deposition in the medial-temporal regions of the brain, vital for episodic memory.^35^ While conflicting evidence on findings related to APOE genotype and tau pathology exists,^36^ the latter may well play a role in the cognitive decline seen in our study. However, in relation to both potential pathological mechanisms,it is unclear whether these are gradual cumulative processes or temporally-specific activations. If the former, we might expect to see these negative associations strengthen with age rather than simply persist.

Episodic memory has been noted in cognitive test batteries to be a highly sensitive marker of incident AD prior to other clinical manifestations.^37,38^ Rates of diagnosed Mild Cognitive Impairment and AD are low in NSHD, in keeping with the comparatively young age of the cohort when considering these diagnoses. It remains possible that the detected associations between APOE-ε4 and episodic memory seen here are attributable to prodromal AD. Given that this decline over almost three decades precedes estimations of the functional manifestations of prodromal AD,^39,40^ continued follow-up of NSHD will determine the age-specific incidence of clinical dementia in those with the APOE-ε4 allele, in comparison to those without. Regardless, the possibility of APOE-ε4 being associated with a process of cognitive decline that begins subtly in midlife, may warrant the search for potential therapeutic window to these deleterious effects, or the instigation of a screening process for potential AD, earlier in the life course.

## Acknowledgements

The NSHD, MJR, DK, MR and AW are supported by core funding and grant funding (programme codes: MC_UU_12019/1, MC_UU_12019/3) from the UK Medical Research Council. DD is funded through a Wellcome Trust Intermediate Clinical Fellowship (WT107467). RB is supported by the National Institute on Aging of the National Institutes of Health under award number P01AG043362. The funders had no role in study design, data collection and analysis, decision to publish, or preparation of the manuscript. Sincere thanks to all of the members of the 1946 birth cohort for their historic and continued participation in NSHD.

## Data and code availability

Data and code are available on request to the NSHD Data Sharing Committee. NSHD data sharing policies and processes meet the requirements and expectations of the UK Medical Research Council (MRC) policy on sharing of data from population and patient cohorts. Data requests should be submitted to; further details can be found at http://www.nshd.mrc.ac.uk/data.aspx. These policies and processes are in place to ensure that the use of data from this national birth cohort study is within the bounds of consent given previously by study members, complies with MRC guidance on ethics and research governance, and meets rigorous MRC data security standards.

